# Optimal Liquid-Based DNA Preservation for DNA Barcoding of Field-Collected Fungal Specimens

**DOI:** 10.1101/2023.11.05.565722

**Authors:** Yu-Ja Lee, Guan Jie Phang, Che-Chih Chen, Jie-Hao Ou, Yin-Tse Huang

## Abstract

Preserving fungal tissue DNA in the field is essential for molecular ecological research, enabling the study of fungal biodiversity and community dynamics. This study systematically compares two liquid-based preservation solutions, RNAlater and DESS, for their effectiveness in maintaining macrofungi DNA integrity during field collection and storage. The research encompasses both controlled experiments and real-world field collections. In the controlled experiments, two fungal species were preserved in RNAlater and DESS at different temperatures and durations. DNA extraction success rates were high, but DNA quality and quantity metrics exhibited variations across samples. However, both preservation solutions demonstrated their viability for preserving fungal DNA, with no significant differences between them. In the field-collected macrofungi experiment, 200 fungal specimens were collected and preserved in RNAlater and DESS. The DNA extraction success rate was 98%, with a few exceptions. The statistical analysis, including paired and independent t-tests, showed no statistically significant differences in DNA quality and quantity between the two preservation methods for the field-collected fungal samples. Overall, this study provides valuable insights into the effectiveness of RNAlater and DESS for preserving macrofungi DNA in field conditions. Researchers can confidently choose between these methods based on their specific needs, without compromising the integrity of the DNA. This research contributes to the advancement of fungal molecular ecology and has broader implications for DNA preservation strategies in ecological and environmental studies.

## Introduction

Preservation of fungal tissue DNA in the field is a crucial aspect of molecular ecological research, enabling the study of fungal biodiversity, community dynamics, and their functional roles in natural ecosystems. However, the challenge lies in effectively preserving fungal tissue DNA in the field, where the samples are exposed to various factors that can degrade DNA integrity. The choice of preservation method, the biological characteristics of the sample, and the duration of exposure to adverse conditions also play critical roles in ensuring successful DNA preservation for subsequent analysis.

Over the years, researchers have made significant efforts in developing preservation methods to maintain the quality and quantity of fungal DNA during collection and transportation. Conventionally, freezing samples at low temperatures (-180 or -20□) has been a common practice to prevent DNA degradation. However, this approach is often impractical in remote field settings where access to freezing facilities is limited. Consequently, alternative preservation methods, such as card-based (FTA® cards, FTA Elute® cards) and liquid-based [DNAgard™, DMSO/EDTA/saturated sodium chloride (DESS), RNAlater®] preservation methods have gained attention for their potential to stabilize nucleic acids under field conditions.

FTA technology employs FTA cards containing cell-lysing and nucleic acid-preserving chemicals (Gray et al., 2013). For extended storage, tissues can be immersed in a high-salt solution like RNAlater®, which has demonstrated superior DNA yield compared to FTA® cards and effectively preserves RNA (Gray et al., 2013). Additionally, a nonproprietary solution DESS has been utilized for DNA preservation. In a comparative study, liquid-based preservatives, including DNAgard™, RNAlater®, and DESS, outperformed card-based methods without one liquid technique showing a clear advantage (Gray et al., 2013).

Prior research has extensively investigated the efficacy of these reagents in safeguarding DNA extracted from a wide range of biological specimens, spanning plant tissues (Kruse et al., 2017; Mbogori et al., 2006), animal tissues (Allen-Hall and McNevin, 2012; Régnier et al., 2011), and the microbial communities residing within host tissues (Carvalhais et al., 2022; Lee et al., 2019; Ndunguru et al., 2005). Nevertheless, while some investigations have delved into their utility in preserving microbe’s, including fungal DNA, there exists a substantial knowledge gap pertaining specifically to the preservation of macrofungi tissues, such as mushrooms. This gap pertains to our comprehension of how these liquid-based preservation methods affect the integrity, quantity, and PCR amplification success of DNA extracted from such macrofungi specimens.

We opted for RNAlater and DESS over other card-based methods due to their superior DNA preservation capabilities (Gray et al., 2013), cost-effectiveness, and ease of specimen transport. This study conducts a systematic comparison of these two preservation reagents, RNAlater and DESS, specifically targeting varied macrofungi tissue DNA. Our experiment encompasses both short-term and long-term preservation scenarios, allowing us to assess DNA quality, quantity, and PCR success rates at multiple time points (day 1 to day 30). By undertaking this analysis, we seek to provide insights into the most effective preservation method for maintaining macrofungi DNA integrity in the field over extended periods. This research not only contributes to the advancement of fungal molecular ecology but also has broader implications for DNA preservation strategies in ecological and environmental studies.

## Materials and Methods

### Sample collection and preservation

We prepared two liquid-based preservation solutions, DESS and RNAlater, following the methods outlined in Seutin et al., (1991) and a patent of Lader, (2012), respectively. For the DESS solution, we added 60 mL of deionized water to 0.25 M (27.9 g) EDTA-2Na (Ethylenediaminetetraacetic acid disodium salt dihydrate, molecular weight 372.24), stirring continuously with a magnetic stirrer, which resulted in noticeable precipitation. Subsequently, we adjusted the pH to 7.5 using 1N NaOH, utilizing approximately 80 mL, which facilitated the gradual dissolution of EDTA-2Na. We then topped up the mixture with deionized water to attain a total volume of 240 mL, before adding 60 mL DMSO (Dimethyl Sulfoxide) and 30 g Sodium Chloride. Thereafter, the solution was autoclaved for sterilization and left to settle overnight at 16□, allowing the excess NaCl to precipitate. The following day, the clear supernatant was meticulously aliquoted into 2 mL tubes, dispensing 1 mL per tube. To ensure sterility, the tubes were subjected to a subsequent round of autoclaving. For RNAlater (5 mM Sodium Citrate, 10 mM EDTA, 70 % ammonium sulfate, pH 5.2), we mixed 40 ml of 0.5 M EDTA, 25 ml of 1 M trisodium citrate, 700 g ammonium sulfate, and 935 ml deionized water, adjusted the pH to 5.2 using 1 M sulfuric acid, and let it cool to room temperature.

In this study, we conducted two experiments: 1) focusing on short- and long-term preservation at 25□ and -20□ (the controlled experiment) and 2) assessing the effectiveness of these methods with field-collected macrofungi samples.

In the controlled experiment, we purchased two commercially available mushrooms (*Pleurotus ostreatus* and *Lentinula edodes*) and preserved their fungal tissues in DESS and RNAlater tubes, storing them at either 25□ or -20□. At designated intervals (day 1, 10, 20, and 30), we extracted DNA from 20 mg of the preserved samples (triplicate per sample) using our in-house DNA extraction workflow outlined below.

For the field-collected macrofungi experiment, conducted from February to July 2023, we engaged citizen scientists to collect fungal samples across Taiwan. These volunteers used collecting tubes containing DESS and RNAlater, dividing each fungal sample into two parts, one placed in DESS and the other in RNAlater. The samples were kept at 4□ or -20□ by citizen scientists and promptly shipped to the laboratory for secure preservation at -20□.

### DNA extraction, PCR, sequence analysis

For DNA extraction, fungal tissues (5–20 mg) underwent lysis with a combination of mechanical beads (1 mm and 0.5 mm zirconia beads in a 1:1 ratio) subjected to agitation at 3200 rpm for 4 minutes using a Vortex-Genie 2 (Scientific Industries, Inc., NY, USA) on its Horizontal Microtube Holder accessory. DNA extraction of lysed cells was carried out using a magnetic-bead-based approach (Phang et al., 2023) adapted from BOMB protocol (Oberacker et al., 2019) on a ZiXpress 32 automated nucleic acid purification instrument (New Taipei City, Taiwan). The quantity and quality of the eluted DNA were assessed using an Agilent BioTek Take3 Multi-Volume Plate following the manufacturer’s instructions. For each metric, triplicate measurements were conducted, and the resulting values were averaged.

For molecular identification, we performed amplification of the entire rDNA regions, encompassing the 18S small subunit (SSU), ITS, and 28S large subunit (LSU). The amplification followed a two-step barcoding procedure as outlined by (Herbold et al., 2015). In the first-step PCR reaction, with a volume of 25 μl, we included 1 μl of DNA template, 0.5 μl of the forward NS1B1ngs_H1 (GCTATGCGCGAGCTGCCCTNGTTGATYCTGCCAGT) primer (including the universal head sequence) adapted from (Tedersoo et al., 2018), 0.5 μl of the reverse LR5_H1 (GCTATGCGCGAGCTGCTCCTGAGGGAAACTTCG) primer (including the universal head sequence) adapted from (Vilgalys and Hester, 1990), 12.5 μl of PowerPol 2X PCR Mix (CAT#RK20718) (Woburn, MA, USA), and 9.5 μl of deionized water. The amplification protocol entailed an initial denaturation at 98□ for 45 seconds, followed by 25 cycles of denaturation at 98°C for 10 seconds, annealing at 64°C for 30 seconds, primer extension at 72□ for 2.5 minutes, and a final extension step at 72°C for 5 minutes. Amplified products underwent purification using DNA Select Isolation BeaverBeads™ (Beaver, China) with a 0.45X product-to-bead volume ratio, following the manufacturer’s protocol. In the second-step PCR reaction, totaling 15 μl in volume, we used 0.75 μl of the first-step purified DNA product as the template, 0.6 μl of barcoded primers, 7.5 μl of 2X ZEJUFast PCR Master Mix (Kaohsiung, Taiwan), and 6.15 μl of deionized water. The second-step PCR amplification protocol entailed an initial denaturation at 98□ for 30 seconds, followed by 12 cycles of denaturation at 98°C for 15 seconds, annealing at 65°C for 15 seconds, primer extension at 72□ for 20 seconds, and a final extension step at 72°C for 3.5 minutes. Subsequently, the barcoded amplicons were sequenced on an Oxford Nanopore Technologies (ONT) MinION sequencer (Oxford, UK) using the manufacturer’s Ligation Sequencing Kit V14 (SQK-LSK114).

The generated ONT sequence data underwent processing through dorado (https://github.com/nanoporetech/dorado) and an in-house compiled pipeline, nanoACT (https://github.com/Raingel/nanoACT). In brief, the process involved basecalling of the ONT raw reads with the model <u>dna_r10.4.1_e8.2_400bps_sup@v4.1.0</u> using dorado; quality filtering with qualityfilt() (QSCORE = 9, MIN_LEN = 400, MAX_LEN = 8000); demultiplexing of barcoded sequences with singlebar(); trimming of artificial sequences such as head and adapter sequences using trim_reads(); de novo clustering of sequences via mmseqs_cluster(); consensus generation of the clustered sequences with mafft_consensus(); and a local BLAST against the NCBI Fungi RefSeq ITS database to obtain preliminary specimen identification through local_blast().

### Statistic analysis

Outliers were identified using the Interquartile Range (IQR) method for metrics, A260/280, A260/230, and DNA concentrations. In short, the bounds for outliers were set at Q1 - 1.5×IQR for lower bound and Q3+1.5×IQR for upper bound. If a sample exceeded these bounds for any metric, this sample and its paired sample were removed from the dataset.

In the exploration of DNA preservation efficacy across distinct conditions, we statistically investigate the impact of various storage temperatures and preservation methods on DNA quality and quantity for two fungal species over multiple days of preservation. The Mann-Whitney U Test, a non-parametric method, was employed to discern statistically significant disparities in DNA metrics (A260/A280, A260/A230, and DNA concentration) amongst groups delineated by the combination of preservation method (DESS and RNAlater) and storage temperature (-20°C and 25°C). This was conducted separately for each day in the preservation timeline, for each species, and for each DNA metric. P-values obtained from the Mann-Whitney U Test provided insights into the days and conditions where significant differences were observed, guiding further in-depth pairwise comparisons using Dunn’s post-hoc test.

To assess the preserving effectiveness of field-collected macrofungi samples preserved in DESS and RNAlater, we carried out paired and independent t-tests on quality metrics, namely A260/A280, A260/A230, and quantity metric, i.e. DNA concentration. The paired t-tests were executed for matched samples within each preserving solution, while the independent t-tests compared the metrics between the two preserving solutions. We visualized the metrics using a boxplot with independent t-test results. In addition, connected scatter plots were generated to illustrate pairwise comparisons of the metrics to show the pattern.

## Results

### Sample collection, preservation, DNA extraction, and PCR

In the controlled experiment, a total of 96 fungal samples were obtained, encompassing two fungal species, two temperature conditions, four-day intervals, two preservation methods, and three replicates for each. DNA of all preserved samples in both DESS and RNAlater preservatives were successfully extracted. Notably, the quality and quantity metrics exhibited variations across samples, with A260/280 ratios ranging from 0.64 to 2.39, A260/230 ratios spanning 0.04 to 2.11, and DNA concentrations ranging from 1.27 to 957.20 (Figure 1).

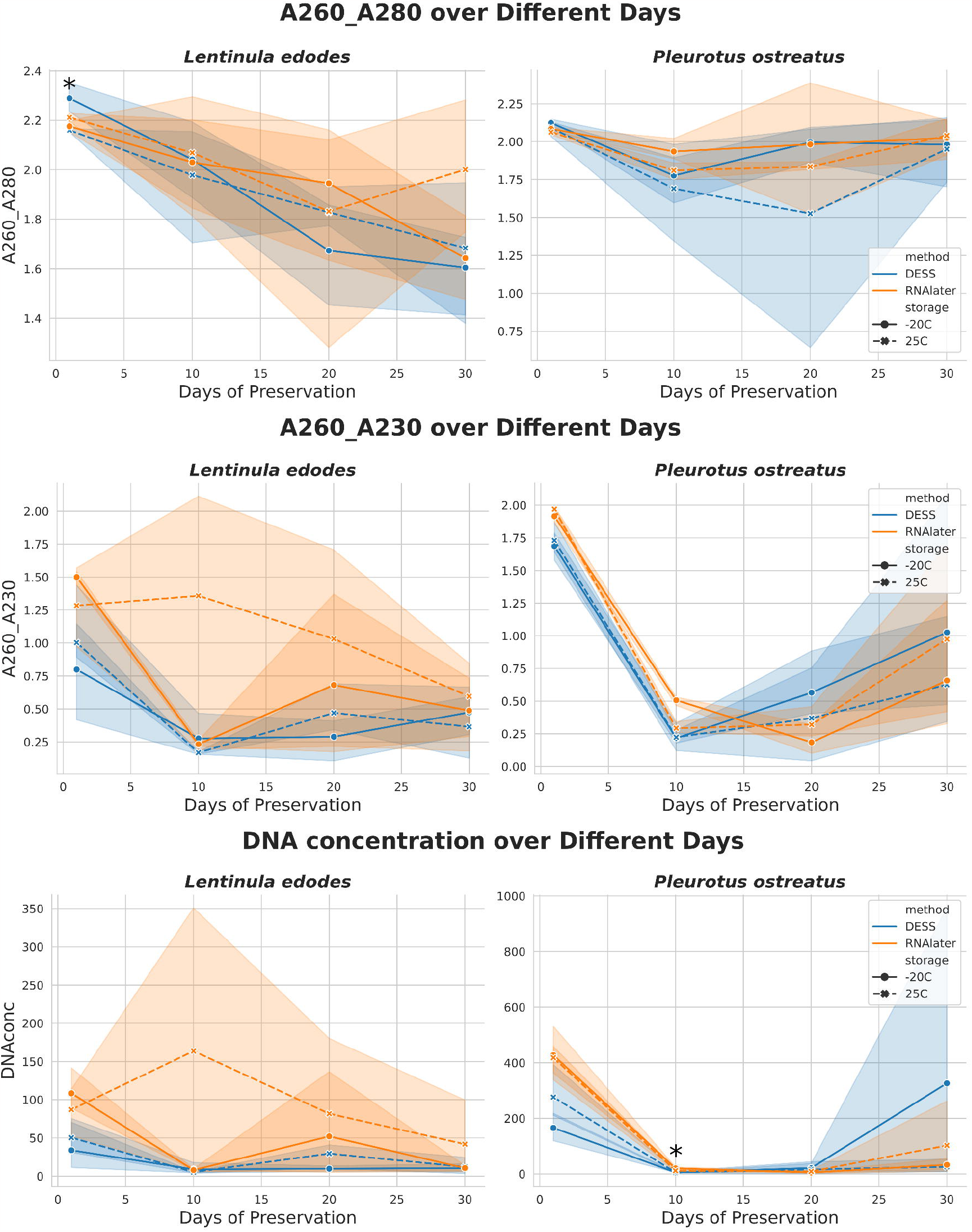

In the field-collected macrofungi experiment, a total of 200 fungal specimens were gathered, comprising 100 fungal isolates preserved in each of the DESS and RNAlater solutions. Among these 100 fungal isolates, there were 67 distinct species identified within the Basidiomycota. DNA extraction success rates reached 98% for both DESS and RNAlater-preserved samples, with the exception of a *Marasmius oreades* and a *Calocera cornea* sample, which were failed in DNA extraction. After removing the outliers, a clean-up dataset consisting of 160 paired samples remained. The quality and quantity metrics displayed variability among the samples, with A260/A280 ratios ranging from 1.45 to 2.59, A260/A230 ratios spanning from 0.10 to 2.28, and DNA concentrations ranging from 1.51 to 265.86. Notably, All DNA-extracted samples in DESS and RNAlater preservatives were successfully amplified for their rDNA regions.

### Statistic analysis

In the controlled experiment, investigating the variation in DNA metrics across different preservation conditions, the Mann-Whitney U Test was employed to discern statistically significant disparities in the DNA quality and quantity metrics (A260/A280, A260/A230, and DNA concentration) across distinct groups, defined by a combination of preservation methods and storage temperatures, for each day of preservation. When comparing preservation methods, DESS and RNAlater showed no significant differences in the A260/A280, A260/A230, and DNAconc measurements, with p-values of 0.214, 0.124, and 0.051 respectively (Table 1). In terms of storage temperature, samples stored at -20°C and 25°C were comparable across all metrics. Comparisons between two different mushroom species, *L. edodes* and *P. ostreatus*, revealed no significant difference in the metrics across the preservation methods, indicating that the effect of preservation might be consistent across these species. Notably, certain conditions and time points revealed significant variations (p value ≤ 0.05) (supplementary Table 1). For instance, for *L. edodes* on day 1, there was a significant difference in the A260/A280 metric with a p-value of 0.03. For *P. ostreatus*, a significant result was observed on day 10 for the DNA concentration with a p-value of 0.03. To pinpoint where these differences lie among the groups, a posthoc Dunn’s test with Bonferroni correction was performed. The results indicated that for *L. edodes* on day 1, using the A260/A280 metric, the DESS stored at -20°C and DESS stored at 25°C had a significant difference with a p-value of 0.03. However, for *P. ostreatus* on day 10 considering the DNA concentration, there was a marginally notable difference between the DESS stored at -20°C and RNAlater stored at -20°C with a p-value of 0.07.

In the assessment of DNA quality and quantity of field-collected macrofungi preserved in RNAlater and DESS, we conducted both paired and independent t-tests for three metrics: A260/A280, A260/A230, and DNA concentration. For the paired t-test, which compares the same sample preserved in both methods, the results yielded p-values of 0.3144, 0.2176, and 0.2629 for A260/A280, A260/A230, and DNA concentration, respectively (Table 1). Similarly, the independent t-test, which evaluates samples across the two preservation methods without pairing, produced p-values of 0.3994, 0.2702, and 0.2872 for the same metrics (Table 2). Given that all the p-values are greater than 0.05, we found no statistically significant difference in DNA preservation quality between the two methods for the field-collected fungal samples.

## Discussion

Preserving fungal tissue DNA in the field is pivotal for advancing molecular ecological research. This study scrutinized the efficacy of two liquid-based preservation solutions, RNAlater and DESS, in safeguarding the integrity, quantity, and PCR amplification success of DNA extracted from macrofungi samples exposed to diverse field conditions. The choice of preservation method, the inherent characteristics of the biological sample, and the duration of exposure to adverse environmental conditions all contribute significantly to the success of DNA preservation for subsequent molecular analysis.

### Effectiveness of RNAlater and DESS for DNA Preservation

Our results, which encompassed two key experiments, provide valuable insights into the effectiveness of RNAlater and DESS as DNA preservation solutions for fungal samples. In the controlled experiment, where we evaluated preservation efficacy over various timeframes and temperature conditions, both RNAlater and DESS demonstrated their viability for preserving fungal DNA. This finding is critical, as it suggests that researchers can confidently choose between these two methods based on their specific needs, without compromising the integrity of the DNA.

Moreover, our field-collected macrofungi experiment extended our investigation to real-world scenarios, mirroring the conditions researchers may encounter during ecological studies. We successfully collected and preserved a substantial number of fungal specimens in both RNAlater and DESS, highlighting the practicality of these preservation methods in field settings. Although the majority of DNA extractions yielded high-quality DNA suitable for downstream analysis, a few exceptions underscore the importance of contingency plans in case of DNA extraction failure. The observed variability in quality and quantity metrics across samples emphasizes the need for careful preservation method selection, particularly when dealing with macrofungi, where DNA characteristics may vary significantly between species (Figure 2).

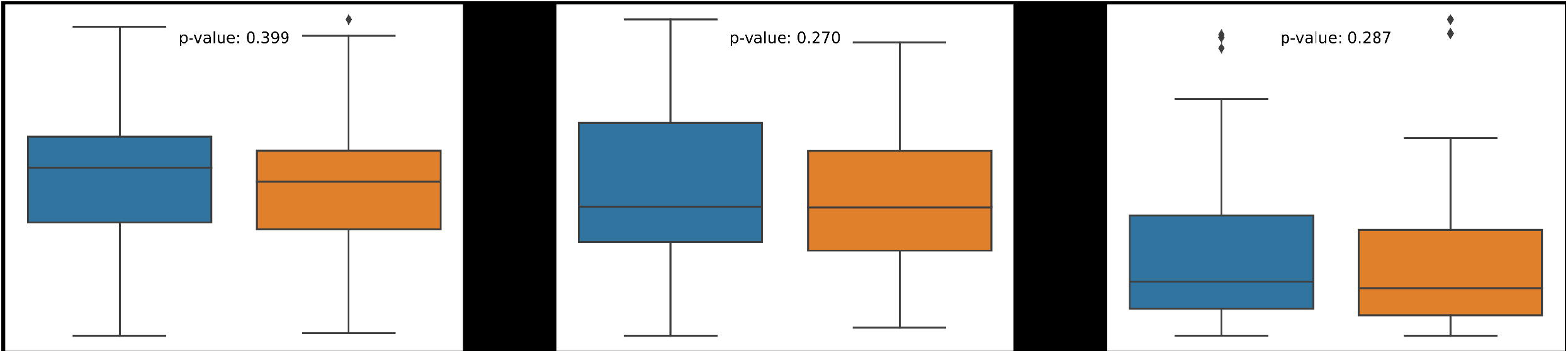

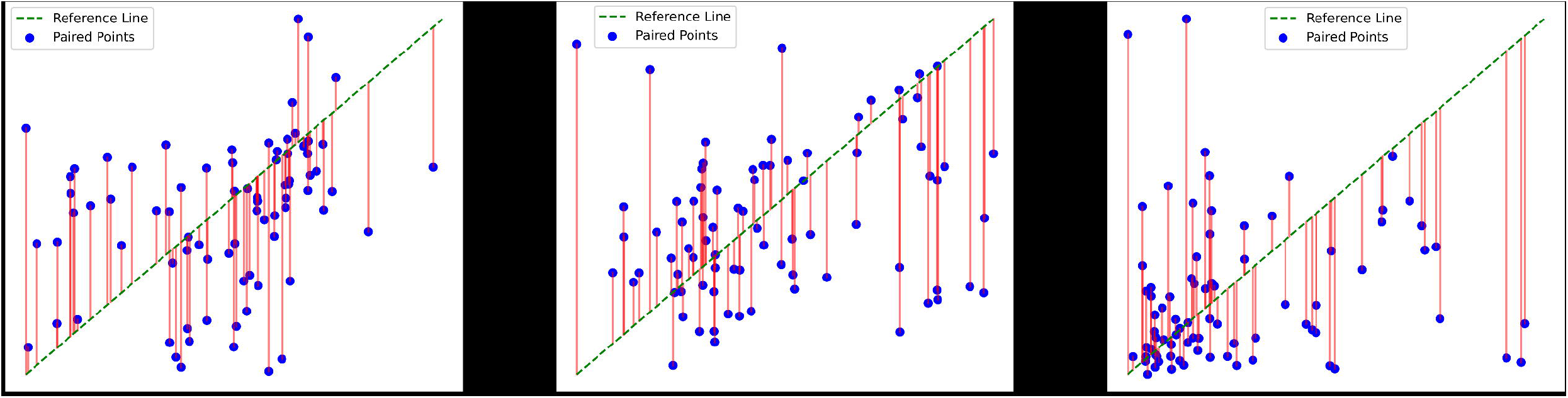

The statistical analysis, employing both paired and independent t-tests, revealed that there were no statistically significant differences in DNA quality and quantity between RNAlater and DESS for the field-collected fungal samples. This finding provides valuable reassurance to researchers seeking cost-effective and reliable DNA preservation methods for their macrofungi studies. The boxplots and connected scatterplots visually illustrate the comparison of DNA metrics between the two preservation methods, further supporting the conclusion of their comparable efficacy.

### Preservation of Tissue Morphology

In addition to preserving nucleic acids, maintaining tissue morphology is crucial for analyzing field-collected samples. However, there’s limited research on this aspect. The impact of RNAlater on tissue morphology remains uncertain. While (Jhavar et al., 2005) achieved excellent HE staining results with RNAlater for human prostate tissue, other studies reported adverse effects on human ovary tissue. DESS appears effective for preserving cartilage morphology over the long term (Staff et al., 2013). Furthermore, DESS preserves nematode adult (Yoder et al., 2006) and egg (Gonzálvez et al., 2022) structures similarly to traditional methods like formaldehyde and low temperatures (Aldeen et al., 1993; Crawley et al., 2016), unlike ethanol, which can distort parasite egg morphology.

Although our study didn’t focus on macrofungi specimen morphology, we observed no significant distortion in spore shape and size. Further research is needed to explore the effects of nucleic acid preservatives on fungal specimen morphology.

## Conclusion

Our study addresses a critical knowledge gap by investigating the effectiveness of RNAlater and DESS as DNA preservation solutions for macrofungi. The results indicate that both preservation methods offer reliable and cost-effective options for maintaining DNA integrity over time. Researchers working with macrofungi in field conditions can confidently choose between RNAlater and DESS based on their specific needs, considering factors such as cost, ease of transport, and logistical constraints. This research contributes to the field of fungal molecular ecology and provides valuable guidance for DNA preservation strategies in ecological and environmental studies.

